## Supplementary information for "Decoupling of timescales reveals sparse convergent CPG network in the adult spinal cord"

*Radosevic et al.*

### Supplementary Information

***Marija Radosevic*[1]*, Alex Willumsen*[1]*, Peter C. Petersen*[1]*, Henrik Lindén*[1]*, Mikkel Vestergaard*[1†] *and Rune W. Berg*[1*]**
[1] *Department of Neuroscience, Faculty of Health and Medical Sciences, University of Copenhagen, Blegdamsvej 3, DK-2200 Copenhagen N, Denmark*
†*Current address: Department of Neuroscience, Max Delbrück Center for Molecular Medicine (MDC), 13125 Berlin-Buch, Germany*

#### Supplementary Discussion

The distribution of slow timescale correlations is qualitatively different for the pairwise intracellular recordings compared with the spike rate correlations of the extracellularly acquired pairs (cf. Fig 5c and 7d). There are likely three reasons for this disparity in the experimental data: 1) The intracellular recordings were directly aimed at identifying relevant rhythmic neurons (Fig. 5b), while not necessarily representing the arrhythmic neurons. 2) The multi-electrode arrays recorded neurons indiscriminately of their relationship with motor activity, and hence the pairwise correlation representing the weakly rhythmic activity cluster around zero. For the same reason we performed the Rayleigh test for rhythmicity to exclude some of the units that were arrhythmic. Nevertheless, many units with weak rhythmicity remained in the population, and these give a peak at zero. 3) The population firing rate distribution is lognormal.[1] Hence, many neurons, even the ones directly related to the rhythm, spike at such a low rate, that the firing rate estimates are poorly modulated by the rhythm. If a pair of neurons - even if their are directly related to the behavior - only fire a couple of time during a motor bout, it is difficult to establish their rhythm and phase. All these pairs make out most of the mass in the distribution around zero in the extracellular data.

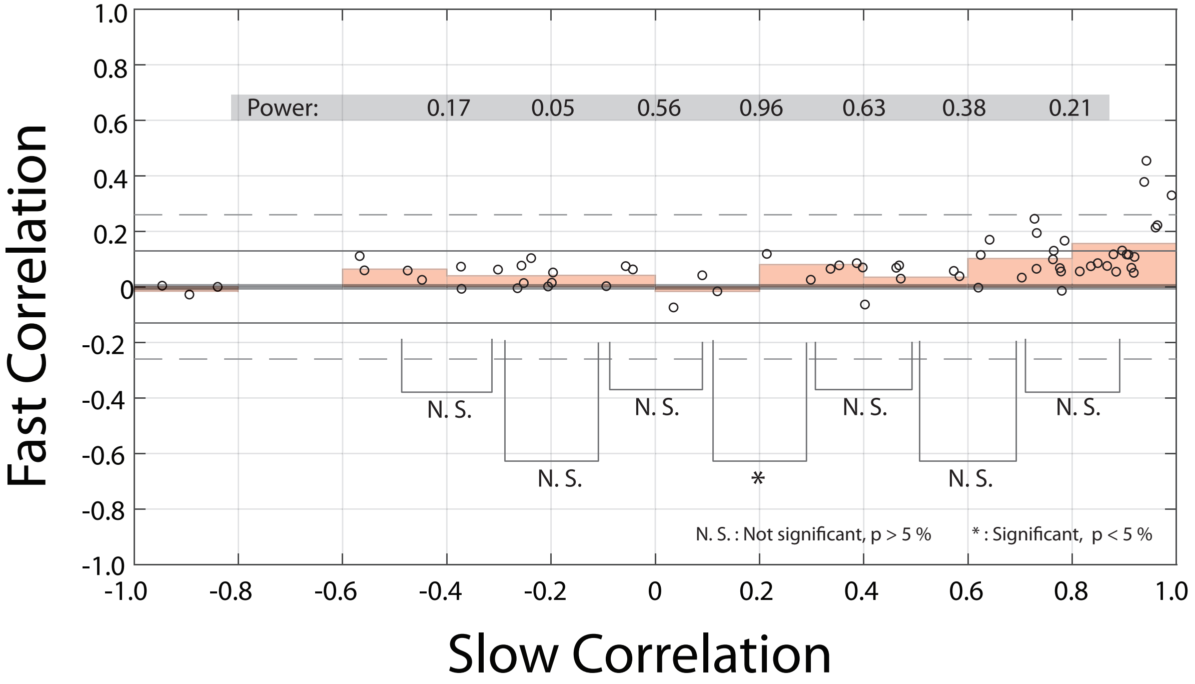


**Supplementary Figure 1:** Pairwise comparisons of bins of slow correlation indicate weak or no significant change. Segregating the slow correlation (Figure 5a in manuscript) into the following bins (red bars): -1.0– -0.8, -0.8– -0.6, -0.6– -0.4, -0.4– -0.2, -0.2–0, 0–0.2, 0.2–0.4, 0.4–0.6, 0.6–0.8, 0.8–1.0. Statistical pairwise comparison of neighboring bins (two–sample t-tests) indicates no significant change at 5% confidence, except one particular case, the bins 0–0.2 and 0.2–0.4 (asterisk symbol). Of particular importance is the comparison 0.6–0.8 with 0.8–1, where the test was unable to reject the null hypothesis of equal mean with 5% confidence. The statistical sample size power for each comparison is indicated above (gray background).
